## Supplementary material for "Engineering indel and substitution variants of diverse and ancient enzymes using Graphical Representation of Ancestral Sequence Predictions (GRASP)"

**Thermal transitions [°C]**

| | $T$m1 | $T$m2 |
|---|---|---|
| *An*GOx | 58.2 | 63.1 |
| N320 | 67.4 | 70.0 |
| N320 Y244E | 71.0 | 73.9 |

Table 1: Comparison of thermal transitions based on differential scanning calorimetry, of an extant glucose oxidase from *Aspergillus niger*, the ancestor inferred at node N320, and the ancestor inferred at node N320 with a single amino acid change, based on marginal distributions.

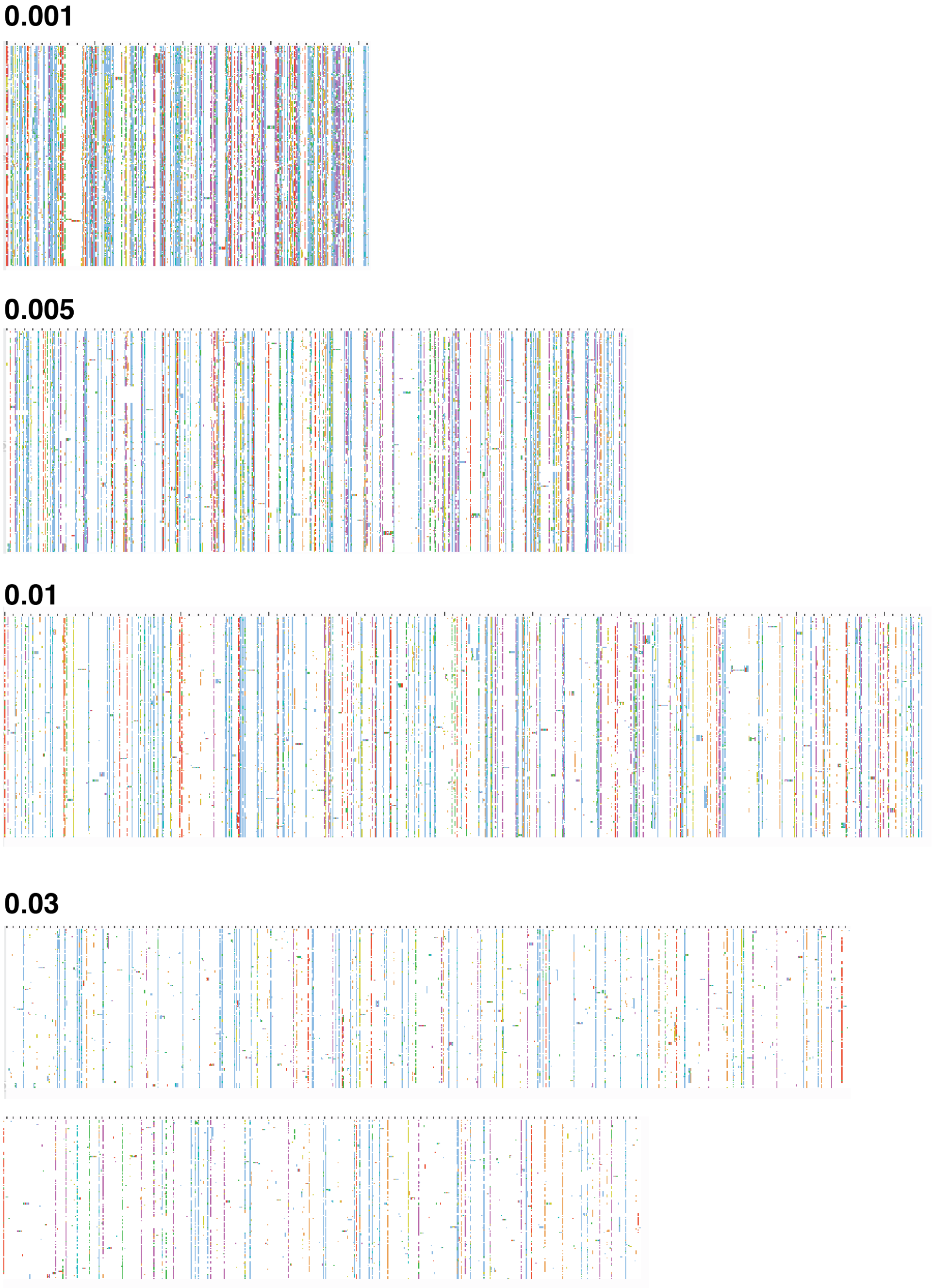


Figure 1: Overview images of the alignments generated for the four indel rates used in the simulated indel evaluation. Each alignment is the true alignment simulated by INDELible for 250 extant sequences with an indel rate of either 0.001, 0.005, 0.01, or 0.03

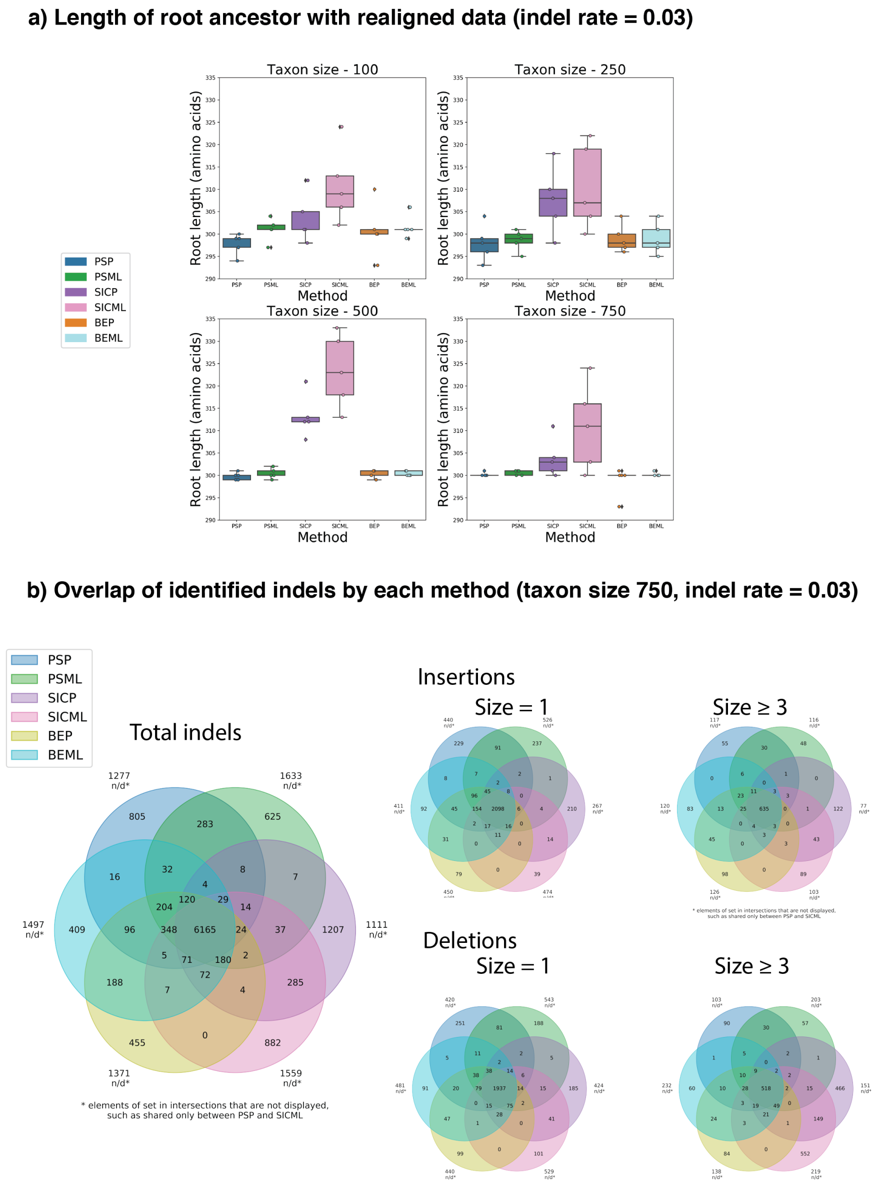


Figure 2: Evaluation of indels with realigned data. **a**, Root length generated from realigned data at four taxon sizes at an indel rate of 0.03 (n=5). **b**, Venn diagram showing the overlap of specific indels identified by each method at taxon size 750 and an indel rate of 0.03. Total numbers of indels are shown in the largest Venn diagram and subsets of this data according to indel type and indel size are shown in the smaller Venn diagrams. Not all intersections are shown ($N = 1$).

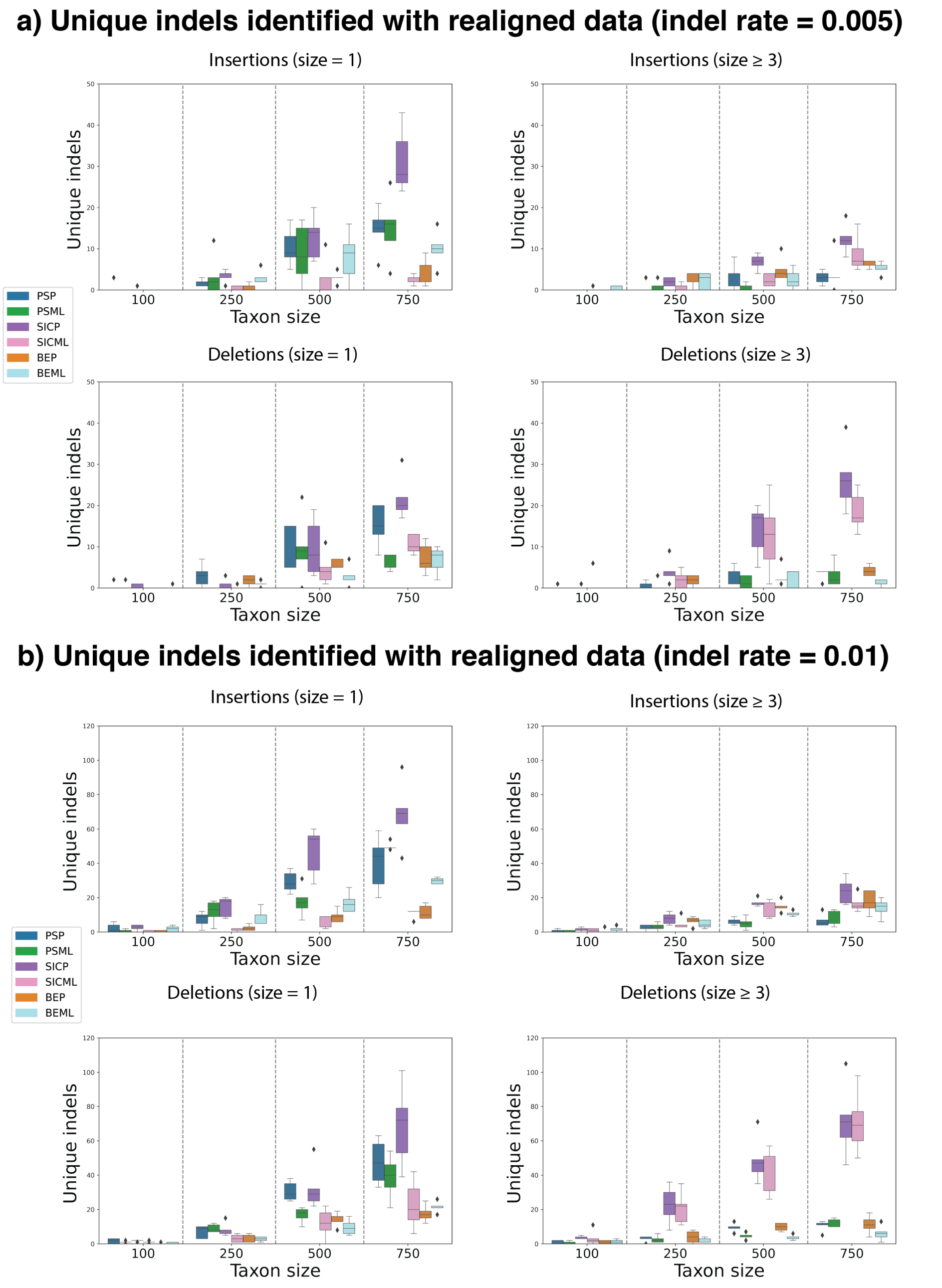


Figure 3: Number of indels uniquely identified by each indel method at four taxon sizes at two indel rates (0.005 and 0.01), organised by indel type and size ($N = 5$). Note the change in range of the y-axis between indel rates.

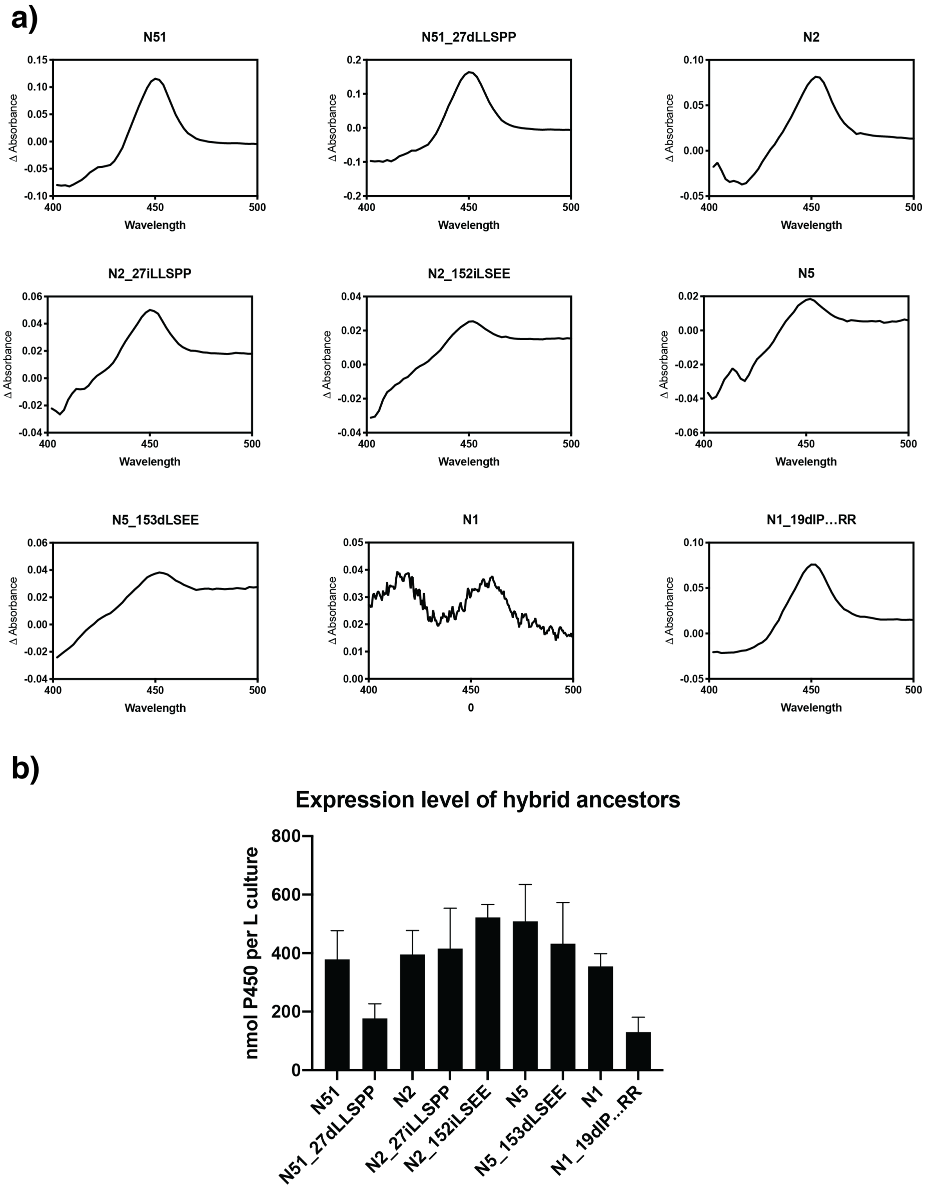


Figure 4: Expression of CYP2U hybrid ancestors. **a**, Fe(II) vs. Fe(II).CO difference spectra for CYP2U ancestors in *E. coli* membranes. **b**, Expression level of CYP2U ancestors in *E. coli* cultures, quantified using Fe(II) vs. Fe(II).CO difference spectroscopy. Data are means +/- SEM, $N = 3$.

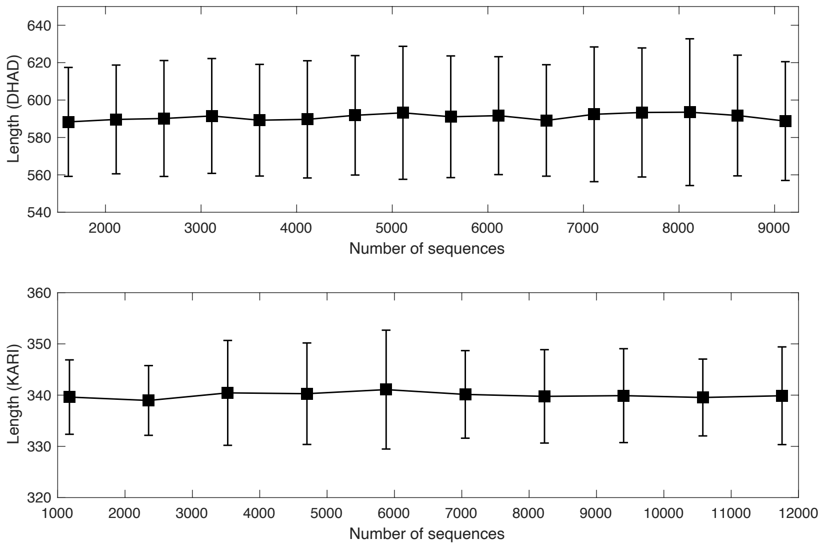


Figure 5: Predicted ancestor sequence lengths are unaffected by size of reconstruction. Mean and standard deviation of the lengths of 50 ancestor sequences mapped are plotted for different reconstructions and data set sizes for DHAD and KARI.

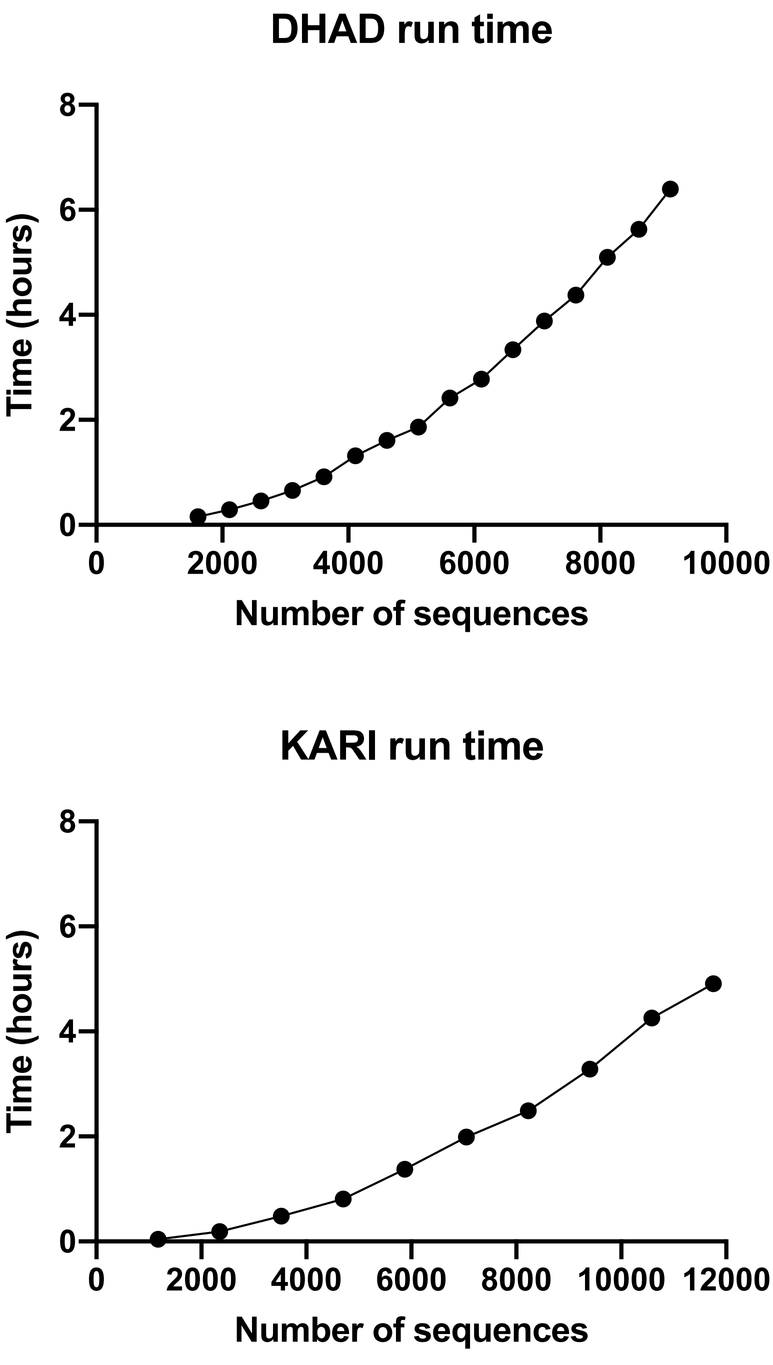


Figure 6: Run times for the DHAD and KARI enzyme families as data set size increases. Reconstructions were performed using GRASP running on 64 GB RAM, 5 threads on 2x 2.6 GHz 14C Xeon VM.

N423 (585 sequence reconstruction)

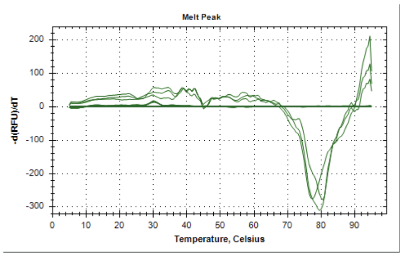


N1442 (9112 sequence reconstruction)

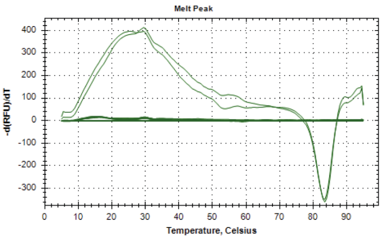


N560 (585 sequence reconstruction)

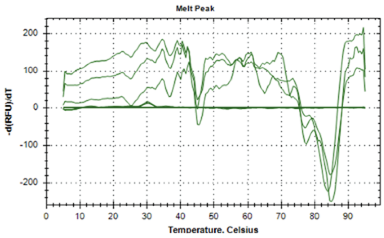


N1443 (9112 sequence reconstruction)

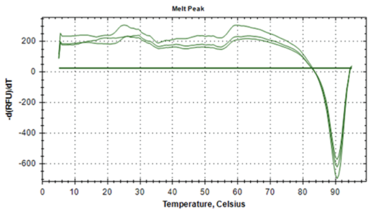


Figure 7: Thermal shift assays showing increase in temperature between equivalent ancestral nodes N423 (585 data set size) and N1442 (9,112 data set size), and equivalent ancestral nodes N560 (585 data set size) and N1443 (9,112 data set size).

a)

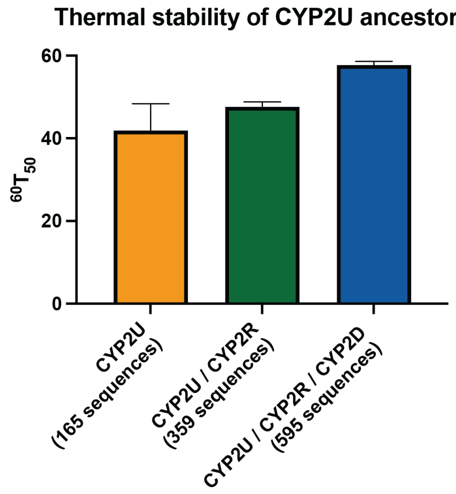


b)

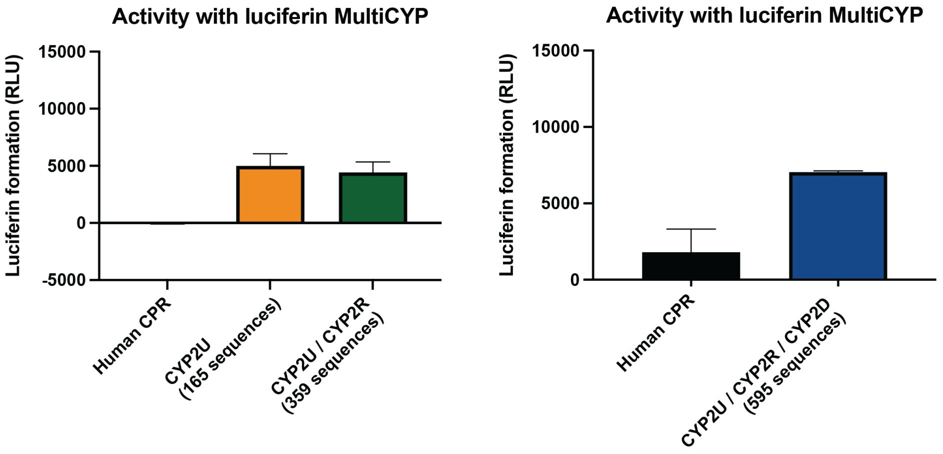


Figure 8: Thermal stability and activity for the CYP2U, CYP2U/CYP2R, and CYP2U/CYP2R/CYP2D ancestors with luciferin MultiCYP. **a**, Comparison of $T_{50}$ values after a 60 minute incubation at a range of temperatures (25-80 °C). Data are means +/- SEM, $N = 2$. **b**, Turnover of luciferin MultiCYP by CYP2U ancestors in *E. coli* membranes, also containing human CPR, after 30 minutes at 37 °C. Membranes from cells expressing only human CPR are included as a negative control. Data are means +/- SEM, $N = 3$. The two graphs represent two independent experiments with two independent levels of human CPR activity recorded.

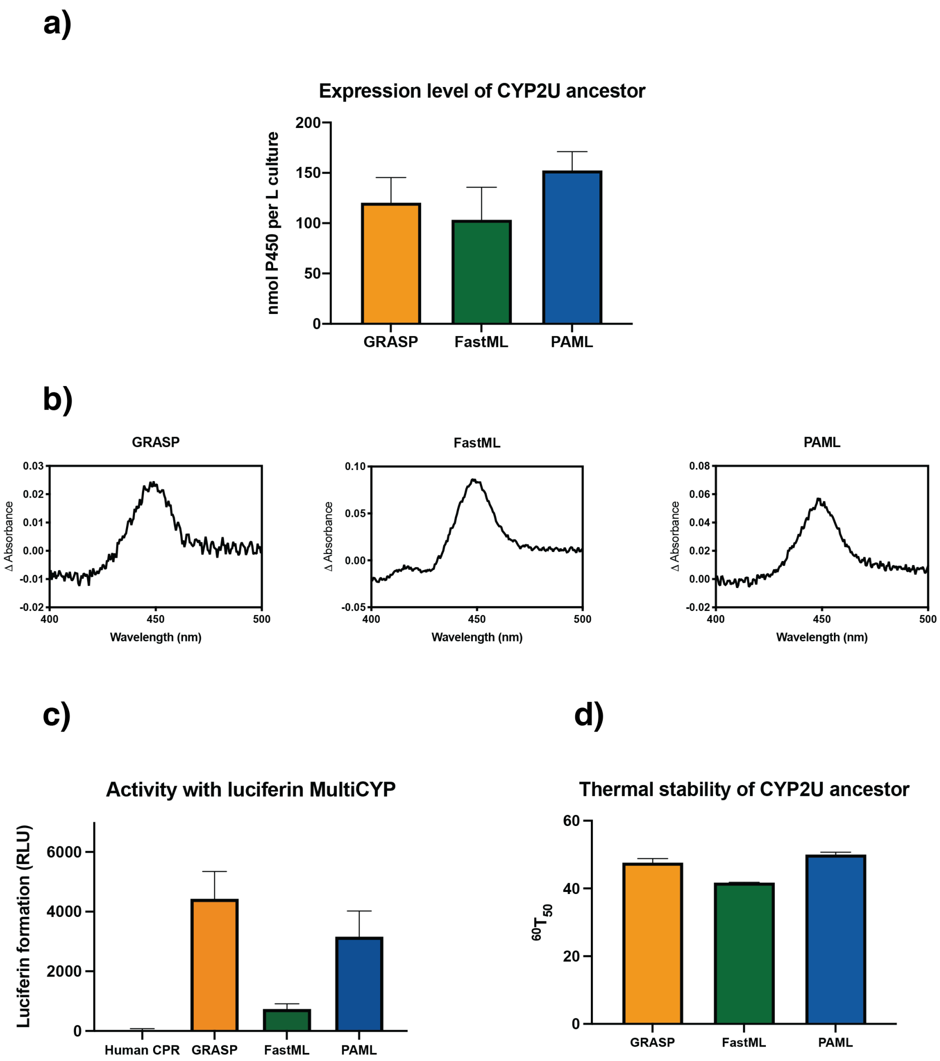


Figure 9: Comparison between ancestors generated using GRASP, FastML, and PAML. **a**, Expression level of CYP2U ancestors in *E. coli* cultures quantified using Fe(II) vs. Fe(II).CO difference spectroscopy. Data are means +/- SEM, $N = 3$. **b**, Fe(II) vs. Fe(II).CO difference spectra for ancestors generated using GRASP, FastML, and PAML in *E. coli* membranes. **c**, Turnover of luciferin MultiCYP by CYP2U ancestors in *E. coli* membranes, also containing human CPR, after 30 minutes at 37 °C. Membranes from cells expressing only human CPR are included as a negative control. Data are means +/- SEM, $N = 3$. **d**, Comparison of $T_{50}$ values after a 60 minute incubation at a range of temperatures (25-80 °C) for ancestors generated using GRASP, FastML, and PAML. Data are means +/- SEM, $N = 2$.

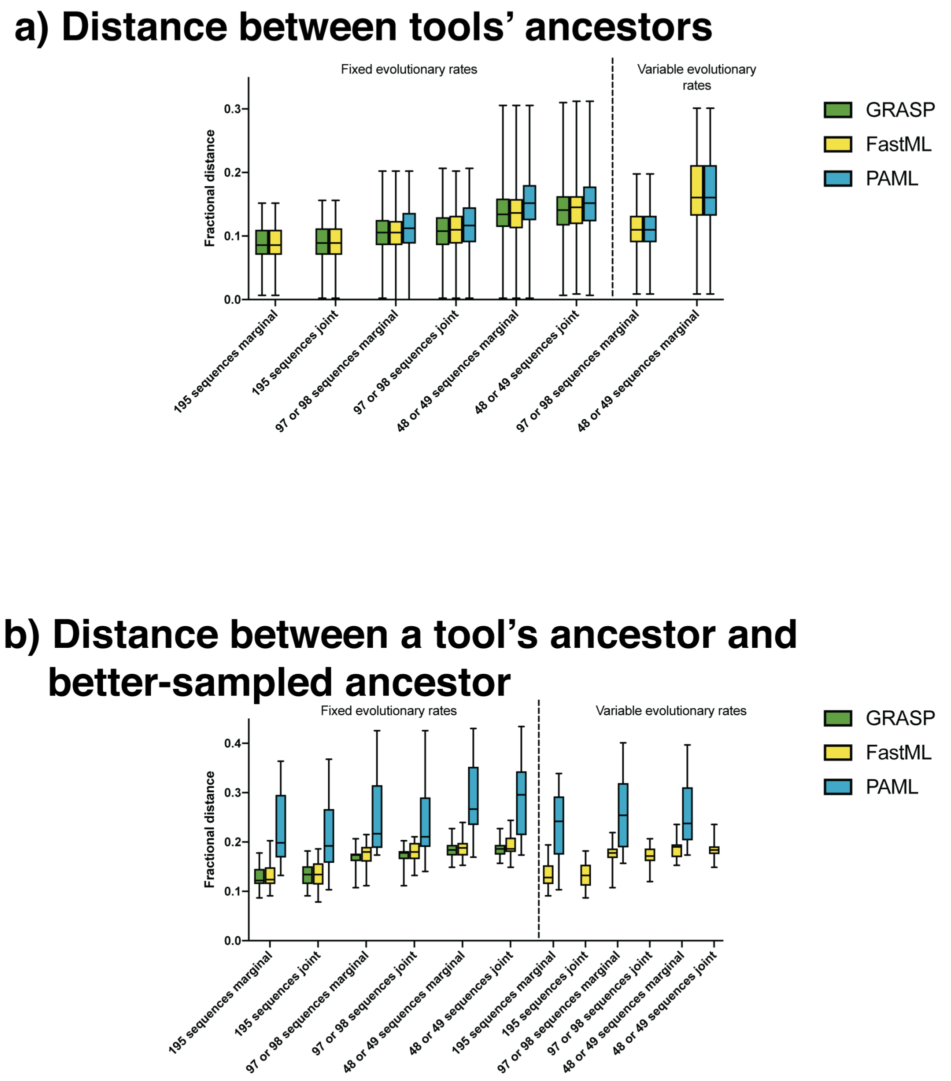


Figure 10: Tool comparison using CYP2 data. **a**, Average fractional distance between tools, calculated as pairwise fractional distances for each ancestral prediction for a given tool against all other ancestral predictions of other tools at 5 groups of 195 sequences, 10 groups of 97 or 98 sequences, and 20 groups of 48 or 49 sequences. Parameter combinations are joint and marginal reconstruction; and fixed or variable evolutionary rates (FastML and PAML only). **b**, Average fractional distance between a better-sampled ancestor inferred by GRASP using 975 sequences and each tool / parameter combination at 5, 10, and 20 groups.

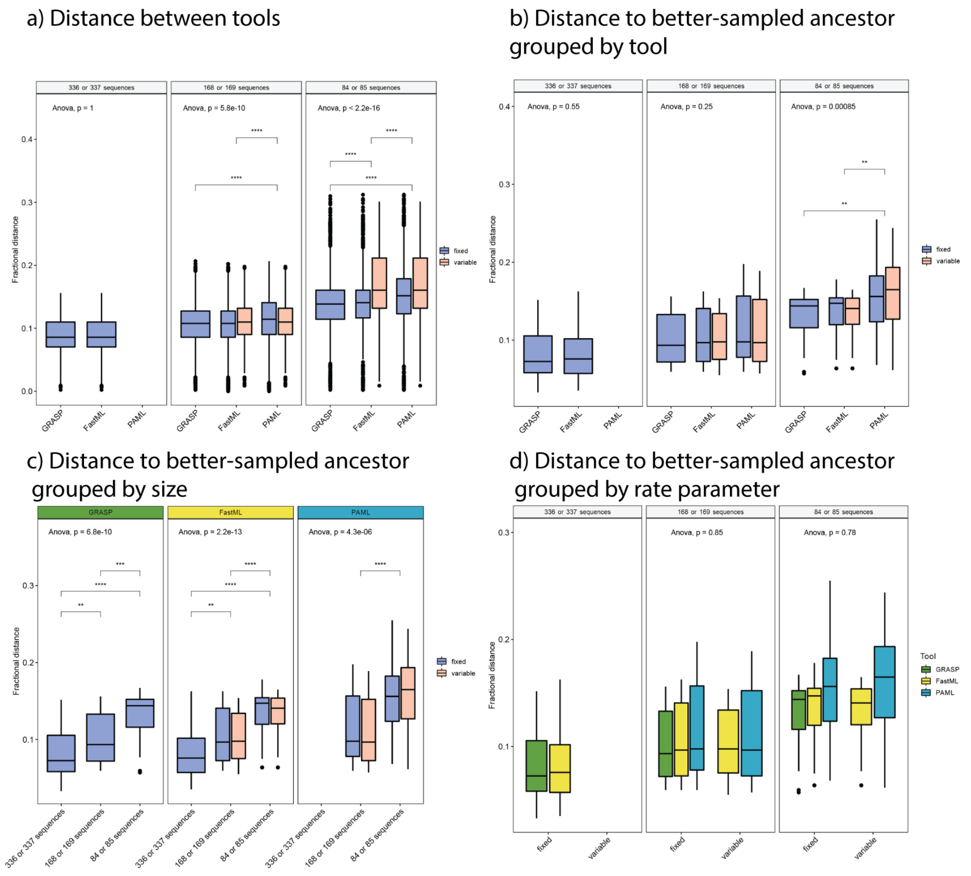

Figure 11: Statistical evaluation of determinants of ancestor prediction performance using 1,682 KARI sequences. **a**, Between tool distances grouped by tool within data set size. **b**, Distance to better-sampled ancestor grouped by tool within data set size. **c**, Distance to better-sampled ancestor grouped by size within tool. **d**, Distance to better-sampled ancestor grouped by rate parameter within data set size. PAML was excluded for the largest data set size; variable rates were not used for the largest data set size. All p-values were determined by a two-tailed Student's $t$-test. Only significant comparisons are shown (* means $p < 0.05$, ** means $p < 0.01$, *** means $p < 0.001$, **** means at limits of precision of test). All parameter settings are from Fig. 5.

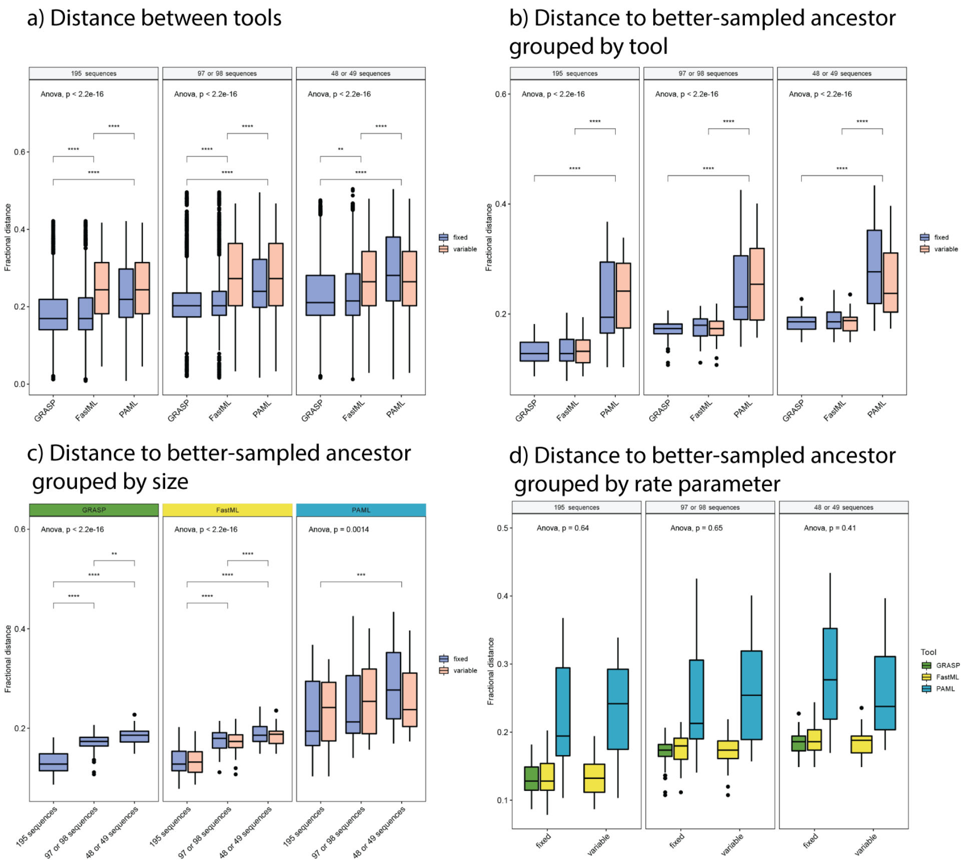


Figure 12: Statistical evaluation of determinants of ancestor prediction performance using 975 CYP2 sequences. **a**, Between tool distances grouped by tool within data set size. **b**, Distance to better-sampled ancestor grouped by tool within data set size. **c**, Distance to better-sampled ancestor grouped by size within tool. **d**, Distance to better-sampled ancestor grouped by rate parameter within data set size. All p-values were determined by a two-tailed Student's $t$-test. Only significant comparisons are shown (* means $p < 0.05$, ** means $p < 0.01$, *** means $p < 0.001$, **** means at limits of precision of test). All parameter settings are from Supplementary Fig. 10.

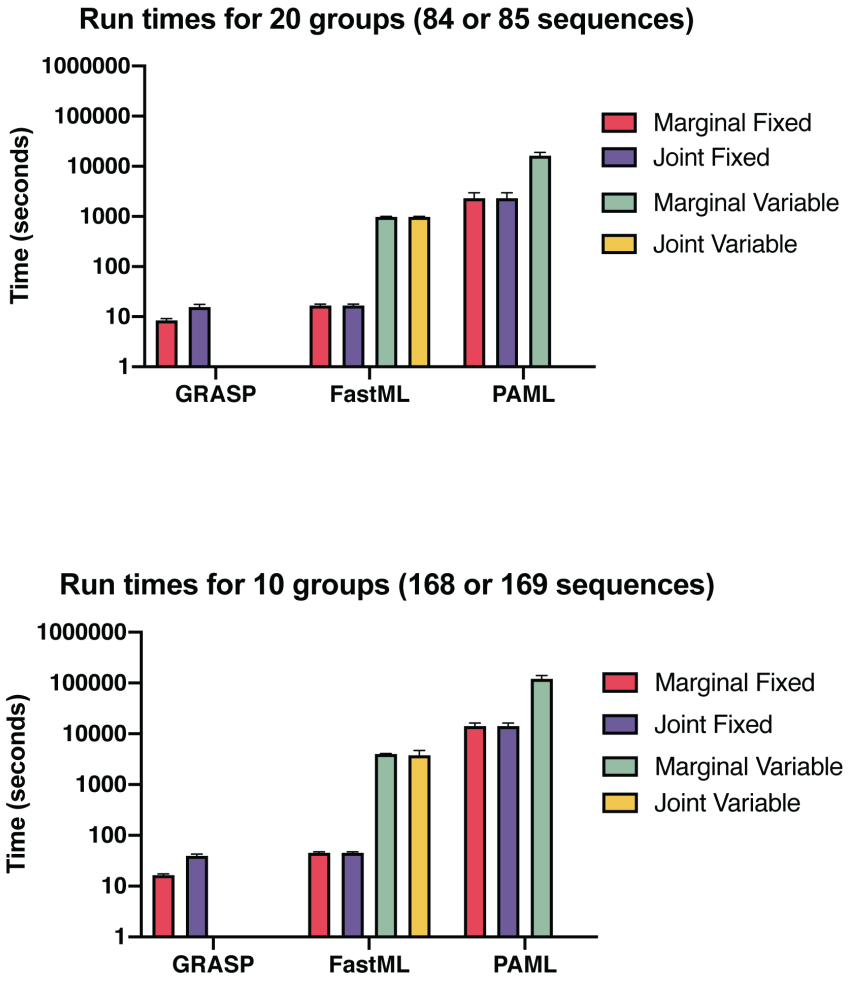


Figure 13: Run times of GRASP, FastML, and PAML at different parameter combinations and group sizes on the KARI data set. Parameter combinations are joint and marginal reconstruction; and fixed or variable evolutionary rates (FastML and PAML only).
